# Tax4Fun2: a R-based tool for the rapid prediction of habitat-specific functional profiles and functional redundancy based on 16S rRNA gene marker gene sequences

**DOI:** 10.1101/490037

**Authors:** Franziska Wemheuer, Jessica A. Taylor, Rolf Daniel, Emma Johnston, Peter Meinicke, Torsten Thomas, Bernd Wemheuer

## Main text

Assessing the functional capability and redundancy of a microbial community is a major challenge in environmental microbiology. To address this challenge, we developed Tax4Fun2, a R-based tool for the rapid prediction of functional profiles and functional redundancy of prokaryotic communities from 16S rRNA gene sequences. By incorporating user-defined, habitat-specific genomic information, the accuracy and robustness of predicted functional profiles can be substantially enhanced.

Microorganisms play a key role in ecosystem functioning^1^. High-throughput sequencing of 16S rRNA genes is a powerful and widely used approach to study the composition and structure of microbial communities in a variety of marine^2–4^, terrestrial^5,6^ and host-associated^7,8^ environments. However, numerous questions in biogeochemistry and ecosystem ecology require knowledge of community functions rather than the taxonomic composition^9^. In recent years, several freely available tools such as PICRUSt^10^, Tax4Fun^11^, Piphillin^12^, Faprotax^13^ and paprica^14^ have been developed. Although these tools cannot replace the functional assessment via metagenomic shotgun sequencing, they have provided unique insights into functional capabilities of prokaryotic communities in diverse habitats, such as soil^5,6^, marine seawater^2,13,14^, microbial mats^15^ and the plant endosphere^7^.

The predictive power of these tools relies on functional information derived from genomes available in public databases. However, available genomes do not necessarily represent the total functional diversity present in the ecosystem investigated. This problem has motivated the development of predictive tools specific for the rumen microbiome^16^ or marine microorganisms^13^. Given the rapidly increasing number of available genomes, in particular through metagenome-assisted genome binning^17^, and that many research groups have access to unpublished, habitat-specific genomic information, the incorporation of this data should enhance the accuracy of functional inferences.

To address these challenges, we developed Tax4Fun2, a novel version of Tax4Fun^11^. Tax4Fun2 is a fast and user-friendly R package (https://sourceforge.net/projects/tax4fun2/) with a current default reference dataset of 275 archaeal and 12,002 bacterial genomes available through NCBI RefSeq database (assessed on 19 August 2018). A novel feature is that Tax4Fun2 can incorporate habitat-specific and user-defined data to increase the robustness and specificity of functional profiles (Fig. 1). Although Tax4Fun2 focuses on prokaryotic data, eukaryotic data can also be incorporated. Tax4Fun2 is platform-independent and highly memory-efficient, enabling researchers without extensive bioinformatics knowledge to predict functional profiles on almost every computer.

**Fig. 1.**
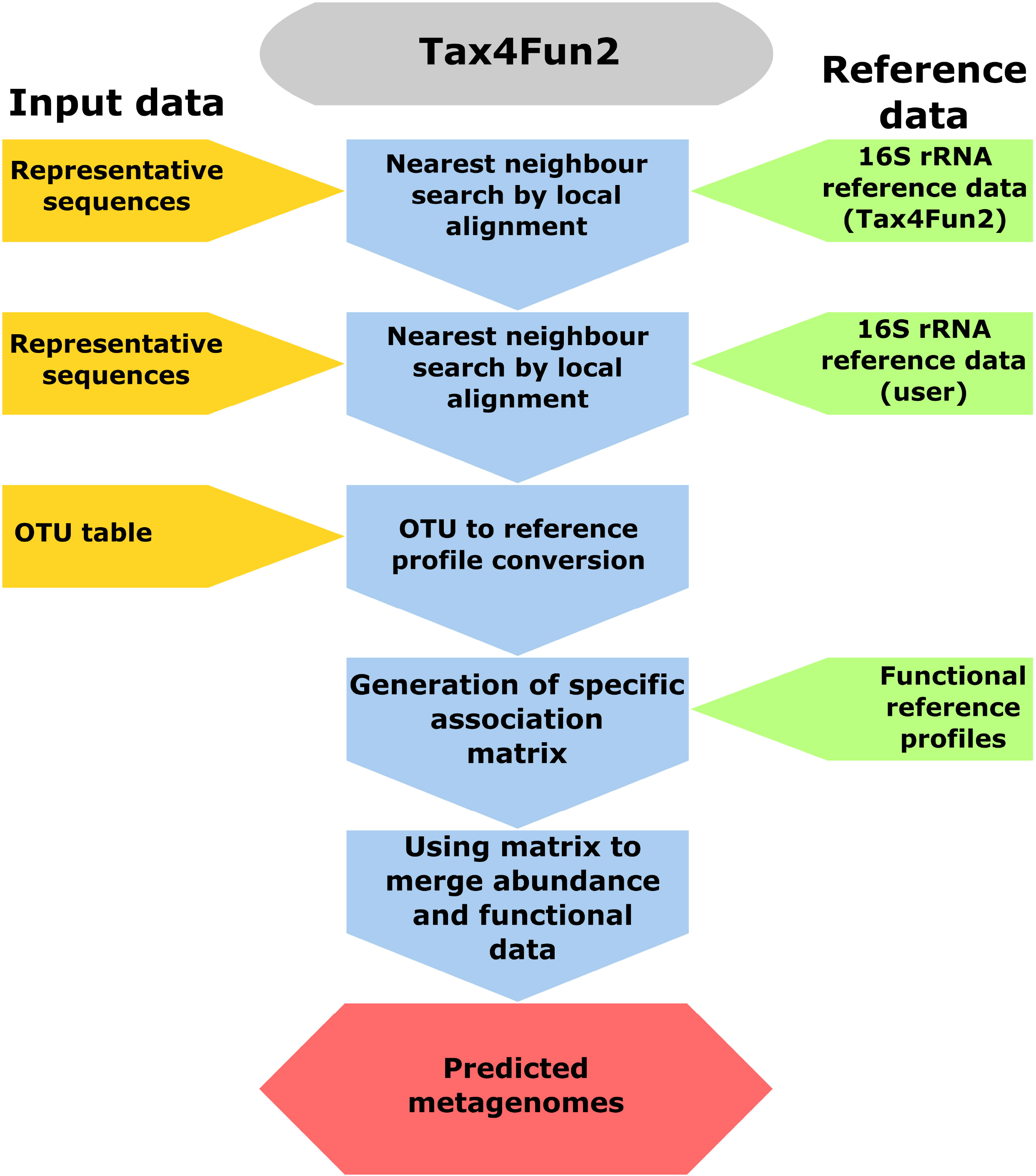
The Tax4Fun workflow. 16S rRNA gene sequences are initially aligned against the reference sequences to identify the nearest neighbour. If user-defined data is supplied, the 16S rRNA gene sequences are additionally aligned against the sequences added by the user. The nearest neighbour in the user data is preferred if both search attempts result in significant hits. The OTU abundances for each sample are summarized based on the results from the nearest neighbour search. An association matrix (AM) containing the functional profiles of those references identified in the 16S rRNA search is generated. The summarized abundances and the functional profiles stored in the AM are merged and a metagenome is predicted for each sample. The amount of sequences/OTUs unused in the prediction is provided in a log file.

We first applied Tax4Fun2 in comparison to Tax4Fun^11^ and PICRUSt^10^ using the same paired samples (16S rRNA and metagenomic data available), which were used to validate Tax4Fun^11^ and PICRUSt^10^, i.e. samples derived from the human microbiome, mammalian guts, soil and from a hypersaline microbial mat (Table 1). In addition, we evaluated the predictive power (defined as high Spearman correlation coefficient) of Tax4Fun2 using ten marine seawater^4^ samples taken in the North Sea and 90 kelp-associated samples collected within the Marine Microbes Framework Data Initiative (http://www.bioplatforms.com/marine-microbes).

**Table 1:**
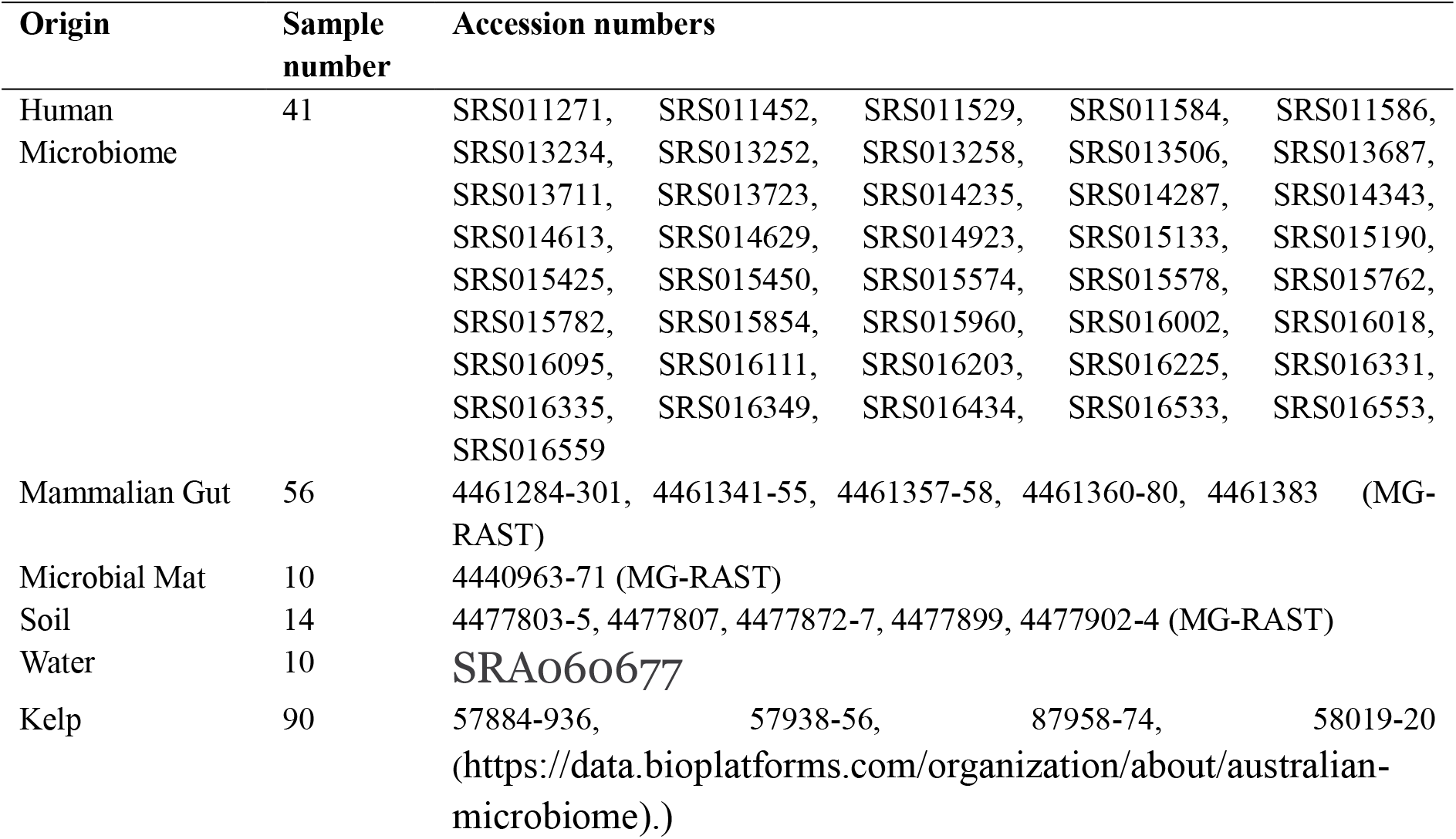
Accession numbers of samples/studies used to validate Tax4Fun2.

Tax4Fun2 outperforms PICRUSt and Tax4Fun across all these datasets (Fig. 2a). Functional profiles predicted by Tax4Fun2 were highly correlated to functional profiles derived from the metagenomes. Although the predicted profiles for the kelp-associated communities were significantly correlated to functional profiles, the median Spearman correlation coefficient was only 0.72, indicating that a lack of suitable reference genomes limits Tax4Fun2’s performance.

**Fig. 2.**
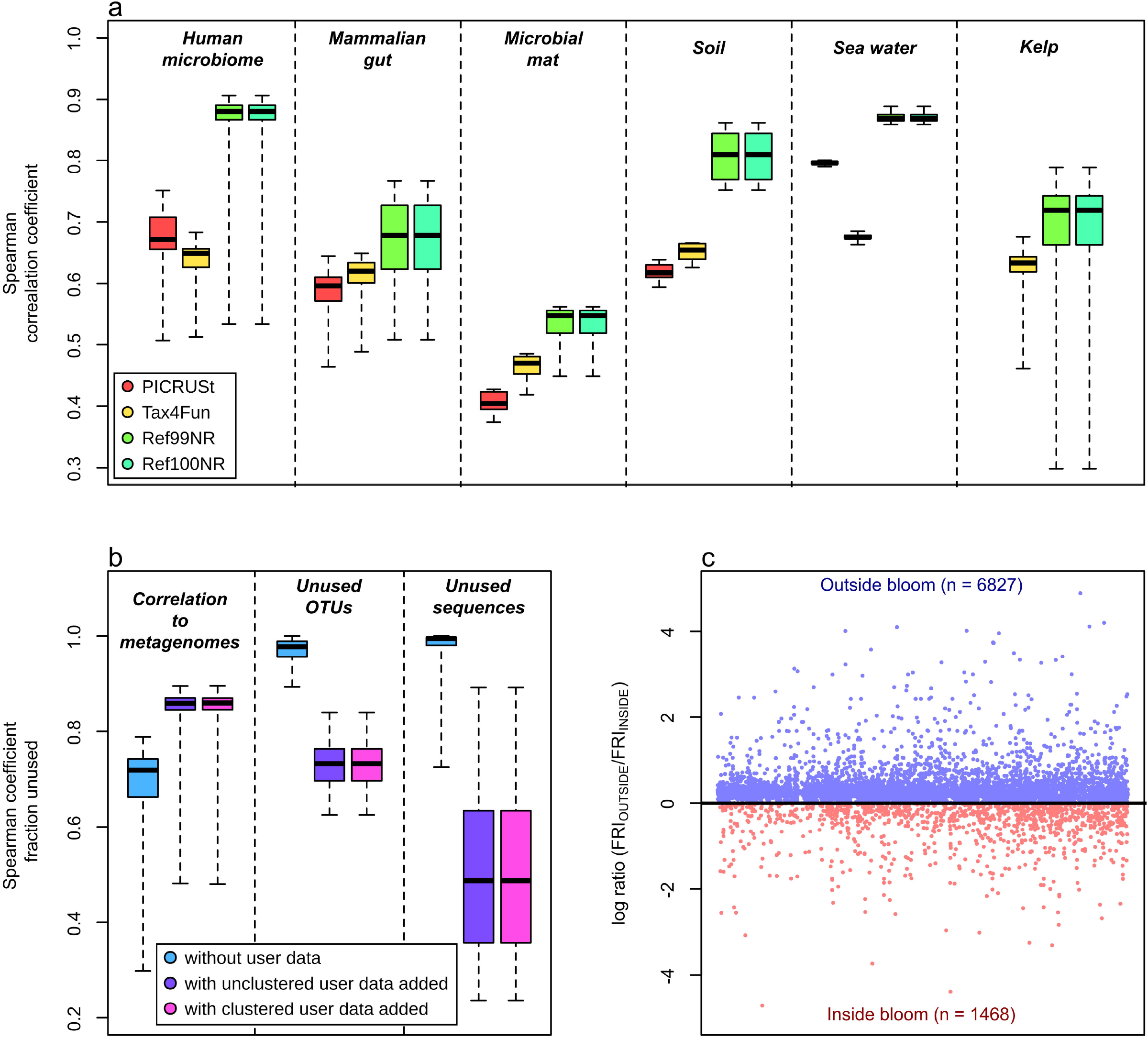
Tax4Fun2 validation. a) Correlations between functional profiles obtained from metagenomic datasets and those predicted from 16s rRNA data. Predictions were made with PICRUSt, Tax4Fun, and Tax4Fun2 using both supplied default reference datasets (Ref99NR and Ref100NR). Note that PICRUSt did not generate any prediction for the Kelp data. b) Correlations between functional profiles retrieved from 90 kelp metagenomes and those predicted with Tax4Fun2 without and with user data added and the fraction of zOTUs and sequences unused in the prediction. C) Functional redundancy indices inside and outside a phytoplankton bloom. A log ratio greater than 0 indicates that a function is more redundant outside the bloom. All predictions were made using a 97% similarity cut off. Correlations are Spearman rank correlations based on relative abundances of KO functions. Only functions present in the metagenome and the predictions were used for comparison. Note that a direct comparison between PICRUSt, Tax4Fun and Tax4Fun2 is difficult due to changes in the KEGG database. Currently, the KEGG databases includes information for more than 10,000 protein-related KO orthologs, whereas PICRUSt and Tax4Fun only provide predictions for around 7000 KO terms.

To address this issue, we used 68 metagenome-assembled genomes (MAGs) derived from the 90 kelp-associated metagenomes to build a kelp-specific genomic dataset. This substantially increased the accuracy of the prediction (median Spearman correlation coefficient with user data added = 0.86) and reduced the fraction of the sequences not used in the predictions (Fig. 2b). Moreover, using the kelp-specific dataset allowed to predict functional profiles for samples, which failed when using only the default reference data because next neighbour search resulted in no close matches. These results demonstrate the benefits of incorporating habitat-specific reference databases, which distinguishes Tax4Fun2 from all other published tools.

A major question in microbial ecology is whether, and to what degree, microbial communities contain functionally redundant members, that may provide stability of ecosystem processes in the face of environmental perturbations^18,19^. In Tax4Fun2, we introduced a functional redundancy index (FRI) with respect to single functions. The FRI is based on the proportion of species capable of performing a particular function and their phylogenetic relationship to each other. A high FRI indicates that a specific function is almost ubiquitous in all community members, whereas a low FRI suggests that the function is present in a few closely related species or has been detected in only one community member. A FRI of 0 indicates that a function is not present at all. Tax4Fun2 calculates a relative FRI (rFRI), which is normalized by the average phylogenetic distance of the community analysed in a specific survey, and the absolute FRI (aFRI), which is normalized by the average phylogenetic distance of all prokaryotes in the reference tree provided with Tax4Fun2. The rFRI can be used to compare samples within one survey, whereas the aFRI allows the comparison of functional redundancy indices across different, unrelated ecosystems.

To test the accuracy of the FRI calculation, we simulated 1,000 communities, each consisting of 100 prokaryotic genomes. We extracted the 16S rRNA gene sequences from each simulated community, clustered them at 97% similarity and calculated the FRI values. We subsequently compared these values to FRI values based on the actual genomic information of the simulated communities. This comparison revealed that Tax4Fun2 provided a good estimate of the functional redundancy present in the microbial community (Spearman rank correlation > 90%) (Fig. 2b). We further tested this approach using the marine sea water samples^4^. Six of these samples were taken inside a phytoplankton bloom and three samples served as reference that were taken outside the bloom. Nearly 7,000 functions displayed a higher functional redundancy in the reference samples, whereas only 1,468 functions had higher redundancies inside the bloom (Fig. 2c). This indicates that the functional redundancy greatly shifts during the phytoplankton bloom. Phytoplankton blooms are usually characterized by a substrate-controlled succession, i.e. distinct bacterial clades dominate the bacterioplankton community at different stages during and shortly after the bloom^20^. Consequently, community members involved in the turnover of certain substrates at a specific stage are predominant and thus their genomes and associated functions will be more redundant, whereas the opposite can be observed for all other community members.

Tax4Fun2 provides researchers with a unique tool to predict and investigate functional profiles of prokaryotic communities based on 16S rRNA gene data. We demonstrated the high predictive power of Tax4Fun2, which can be further enhanced by the incorporation of user-defined and habitat-specific data. Another unique feature of Tax4Fun2 is that it enables researchers to calculate the redundancy of specific functions, which is critical for the prediction how likely a specific function is lost during environmental perturbation. Tax4Fun2 with its user-friendly, simplified workflow will assist researchers considerably in the functional analysis of microbial communities.

## Methods

### Datasets used in this study

To compare Tax4Fun2 with Tax4Fun^11^ and PICRUSt^10^, we used the same 16S rRNA datasets which were originally used to validate Tax4Fun and PICRUSt (for details see ^11^). We further assessed the accuracy of Tax4Fun2 using 10 marine water (taken from ^4^) and 90 kelp-derived metagenomes (for details see https://data.bioplatforms.com/organization/about/australian-microbiome). A list with all accession number is provided in Table 1.

### Processing of 16S rRNA gene data from marine water samples

Pyrosequencing data were processed using QIIME version 1.8^21^. After raw data extraction, reads shorter than 600 bp or longer than 900 bp, exhibiting low quality (<25), possessing long homopolymer stretches (>8 bp), or showing primer mismatches (>2 bp) were removed. Remaining reverse primer sequences were truncated employing cutadapt version 1.18^22^. Processed sequences of all samples were concatenated and denoised employing Acacia version 1.53b^23^. Denoised sequences were sorted by decreasing length and clustered at 97% sequence identity in operational taxonomic units (OTUs) employing the UCLUST algorithm implemented in USEARCH version 8.1.1861^24^. Chimeric sequences were removed using UCHIME^25^ implemented in USEARCH in *denovo* and *reference* mode with the SILVA database (SILVA_132_SSURef_Nr99) as reference dataset^26^.

### Processing of 16S rRNA gene data derived from kelp samples

Paired reads were merged with Flash^27^ and subsequently processed with USEARCH version 10.240^24^. Merged reads were quality-filtered; the filtering included the removal of low-quality reads (maximum number of expected errors >2 and more than 1 ambiguous base) and those shorter than 400 bp. Processed sequences of all samples were concatenated into one file, dereplicated, and obtained unique sequences were denoised and clustered into zero-radius OTUs (zOTUs) with the *unoise3* algorithm. A *de novo* chimera removal was included in the unoise step. Afterwards, remaining chimeric sequences were removed using the *uchime2* algorithm^25^ in high confidence mode with the SILVA database as reference dataset^26^. Subsequently, processed sequences were mapped onto zOTU sequences to calculate the presence and abundance of each zOTU in every sample using the *otutab* command with *maxrejects* and *maxaccepts* options disabled.

### Functional predictions based on 16S rRNA data

Functional profiles were predicted with PICRUSt^10^, Tax4Fun^11^ and Tax4Fun2. For PICRUSt, processed sequences were clustered using QIIME^21,28^ by close reference picking against greengenes (13_5), PICRUSt’s default database. OTU abundances were normalized by 16S rRNA copy numbers prior to the calculation of functional profiles. For Tax4Fun, OTU sequences were aligned against the SILVA database (SILVA_123_SSURef_Nr99) ^26^ using BLAST version 2.7.1^29^. The OTU table and the taxonomic classification were subsequently merged and used to predict functional profiles in Tax4Fun using default settings. For Tax4Fun2, functional profiles were initially aligned against the supplied 16S rRNA reference sequences by BLAST using the *runRefBlast* function. Functional predictions were subsequently calculated using the *makeFunctionalPrediction* function.

### Generation of reference datasets

Tax4Fun2 is supplied with two reference datasets (Ref99NR and Ref100NR) refereeing to the similarity threshold used during 16S rRNA clustering. Each dataset consists of an association matrix with 16S rRNA reference sequences associated with functional reference profiles (number of entries in the association matrix: 4,584 and 18,479 for Ref99NR and Ref100NR, respectively).

Reference datasets were generated as follows: we downloaded all complete genomes and all genomes with the status ‘chromosome’ from NCBI RefSeq (assessed on 18 August 2018), resulting in 275 archaeal genomes and 12,102 bacterial genomes. Barrnap version 0.9 (https://github.com/tseemann/barrnap) was used to identify and extract all 16S rRNA gene sequences. All obtained sequences were subsequently concatenated into a single file, sorted by decreasing length and clustered using the UCLUST algorithm implemented in USEARCH version 10.240^24^ at 99% and 100% sequence similarity, respectively. The longest sequence of each cluster served as 16S rRNA reference sequence.

Functional profiles were generated for each genome as follows: open-reading-frames were identified with prodigal version 2.6.3^30^. Functional profiles were calculated based on deduced protein sequences with UProC version 1.2.0^31^ using the KEGG database for prokaryotes (July 2018 release) as reference^32^. To account for differences in rRNA copy numbers, functional profiles were normalized by the number of 16S rRNA genes identified in each genome. Due to the heterogeneity of 16S rRNA genes within a genome, the functional reference profile for each 16S rRNA reference sequence was generated based on the 16S rRNA clustering results: a single functional reference profile is the average normalized functional profiles of each genome with at least one 16S rRNA gene affiliated to the cluster. If more than one 16S rRNA gene sequence of a genome was assigned to the cluster, the normalized profile of the genome was multiplied by the number of 16S rRNA genes before calculating the mean profile.

The algorithm which was used to generate the reference data is implemented in the Tax4Fun2 package (function = *addUserDataByClustering*). Note that a 32-bit version of USEARCH is required to use this function. USEARCH is freely available at https://www.drive5.com/usearch/.

### Testing the predictive power of Tax4Fun2

To test the predictive power of Tax4Fun2 compared to PICRUSt and Tax4Fun, we used the same paired samples (16S rRNA and metagenomic data), which were originally used to validate Tax4Fun’s accuracy. Functional profiles for each metagenome used in the validation process were generated as follow: protein sequences were extracted with prodigal version 2.6.3^30^ and functional annotations were made with UProC version 1.2.0^31^ as described above for the functional genome annotation. We validated the accuracy of PICRUSt, Tax4Fun and Tax4Fun2 by comparing the functional profiles predicted to the metagenomic profile using Spearman correlation (see Figure 2). Due to several changes in the KEGG orthology since PICRUSt and Tax4Fun were developed (deprecated and new functional orthologs), a direct comparison of functional profiles predicted with all three tools is difficult. Hence, functional profiles were converted to relative abundances prior to comparison. Only functions present in the metagenomic profile and in the predictions were considered in the comparison.

### Generation of metagenome-assembled genomes (MAGs)

The incorporation of user-derived genomes is a key feature of Tax4Fun2, allowing users to build their own reference data. To exploit the accuracy of Tax4Fun2 with default settings (without user data) and with a user-defined reference database, we added 68 MAGs obtained from the 90 kelp metagenomes. The genomes were extracted from the metagenomes as follows: raw data were quality trimmed with Trimmomatic version 0.36^33^ and subsequently assembled with metaSPAdes version 3.11.1^34^. The coverage of each scaffold was determined by mapping the processed data on the assembled scaffolds using bowtie version 2.3.2^35^. Scaffolds smaller than 2,500 bp were removed. After converting to bam format and sorting using samtools ^36^, the coverage was determined with the *jgi_summarize_bam_contig_depths* script. Genomes were extracted using MetaBAT version 0.32.5^37^ and MyCC^38^ and subsequently refined using the binning_refiner version 1.2^39^. 16S rRNA gene sequences were identified using barrnap version 0.9. The completeness and contamination was determined with checkM version 1.0.7^40^. All genomes with more than 50% completeness, less than 5% contamination and possessing at least one 16S rRNA gene were included as user data in the Tax4Fun2 prediction. A functional profile was generated with UProC version 1.2.0^31^ and KEGG as described above for the functional genome annotation.

### Calculation of the functional redundancy index (FRI)

In Tax4Fun2, we introduced the functional redundancy index (FRI). The FRI describes the redundancy of any given function in the investigated community. It incorporates the phylogenetic distribution (distance) of community members harbouring the function and their proportion in the community (see Formula 1).

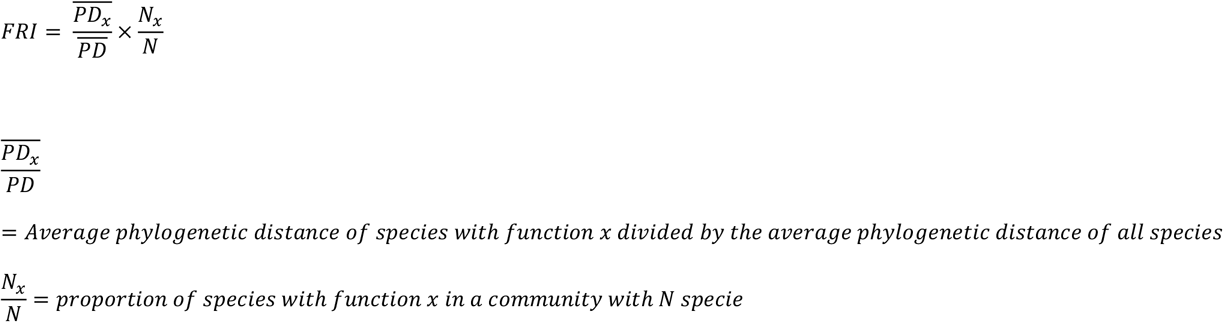

To account for differences in phylogenetic distance, we introduced the absolute and relative FRI (aFRI and rFRI). The difference between them is the average phylogenetic distance used for its normalization. To calculate the aFRI, the average phylogenetic distance of all species in the full 16S rRNA reference tree is used, whereas the rFRI is normalized by the average phylogenetic distance of those species in the 16S rRNA reference tree observed in a sample. The tree for each of the two reference datasets was generated as follows: all 16S rRNA reference sequences were aligned with SINA version 1.2.11^41^ and the latest Silva ARB release (SILVA_132_SSUREF_NR99). The phylogenetic tree was calculated using RaxML version 8.2.11^42^ under a GTRGAMMA model and a random seed of 12345.

### Testing the functional redundancy index using simulated datasets

To test the FRI accuracy, we simulated 1,000 communities each consisting of 100 genomes randomly selected from the 12,377 genomes used to generate the reference data. The genomes were selected based on random numbers generated with the *sample* function in R. To assess the phylogenetic distance between the genomes, we extracted 59 marker protein sequences based on hmm profiles derived from PFAM version 31^43^ and TIGRFAM version 15^44^. The 59 marker proteins were selected because their corresponding genes were present in 90% of all 12,377 genomes and, if present, were single-copy genes in 99% of them. These criteria were applied to archaea and bacteria independently. The extracted protein sequences of each hmm profile were aligned using mafft version 7.3.11^45^. Afterwards, aligned protein sequences for each genome were concatenated. The phylogenomic tree was calculated under a WAGGAMMA model using FastTree version 2.1.10^46^. An initial attempt to use RaxML^42^ failed due to the size of the alignment. Functional profiles for each genome were converted to presence absence data and the FRI was calculated for each function using the genome tree and the presence-absence data. To calculate the FRI in Tax4Fun2, the 16S rRNA gene sequences present in the 100 genomes of each subset were clustered in operational taxonomic units (OTUs) at 97% similarity with UCLUST implemented in USEARCH version 10.240. The longest sequence of each cluster was used to represent each OTU. The FRI was subsequently calculated using Tax4Fun2. The FRIs calculated for each function by Tax4Fun2 were compared to the FRIs calculated directly from the genomes of each simulation by Spearman rank correlation in R. The OTU table necessary for the calculation contained the id and size of each OTU.

## Statistical data analysis

All statistical tests were performed in R version 3.5.1^47^.

## Data availability

Tax4Fun2 is feely available at https://sourceforge.net/projects/tax4fun2/.

## Acknowledgements

This research was partly funded by the German Research Foundation (DFG): research fellowship granted to B.W. and TRR51 granted to R.D. T.T. and J.A.T. were supported by Bioplatforms Australia. F.W. and E.L.J. were supported by ARC Linkage Project SHRP021212.

## Author Contributions

B.W. led the project. B.W., P.M. and F.W. designed and implemented the final Tax4Fun2 algorithms, to which T.T. and R.D. made critical contributions. J.A.T. collected and analysed the kelp dataset used for the validation of Tax4Fun2. B.W., P.M. and R.D. coordinated the online implementation. F.W. and B.W. wrote the manuscript, with feedback from all other authors. All authors approved the final version of the manuscript

